# Flavivirus NS1-triggered endothelial dysfunction promotes virus dissemination

**DOI:** 10.1101/2024.11.29.625931

**Authors:** Henry Puerta-Guardo, Scott B. Biering, Bryan Castillo-Rojas, Michael J. DiBiasio-White, Nicholas TN Lo, Diego A. Espinosa, Colin M. Warnes, Chunling Wang, Thu Cao, Dustin R. Glasner, P. Robert Beatty, Richard J. Kuhn, Eva Harris

## Abstract

The *Flaviviridae* are a family of viruses that include the important arthropod-borne human pathogens dengue virus (DENV), West Nile virus, Zika virus, Japanese encephalitis virus, and yellow fever virus. Flavivirus nonstructural protein 1 (NS1) is essential for virus replication but is also secreted from virus-infected cells. Extracellular NS1 acts as a virulence factor during flavivirus infection in multiple ways, including triggering endothelial dysfunction and vascular leak via interaction with endothelial cells. While the role of NS1 in inducing vascular leak and exacerbating pathogenesis is well appreciated, if and how NS1-triggered endothelial dysfunction promotes virus infection remains obscure. Flaviviruses have a common need to disseminate from circulation into specific tissues where virus-permissive cells reside. Tissue-specific dissemination is associated with disease manifestations of a given flavivirus, but mechanisms dictating virus dissemination are unclear. Here we show that NS1-mediated endothelial dysfunction promotes virus dissemination *in vitro* and *in vivo*. In mouse models of DENV infection, we show that anti-NS1 antibodies decrease virus dissemination, while the addition of exogenous NS1 promotes virus dissemination. Using an *in vitro* system, we show that NS1 promotes virus dissemination in two distinct ways: (1) promoting crossing of barriers and (2) increasing infectivity of target cells in a tissue- and virus-specific manner. The capacity of NS1 to modulate infectivity correlates with a physical association between virions and NS1, suggesting a potential NS1-virion interaction. Taken together, our study indicates that flavivirus NS1 promotes virus dissemination across endothelial barriers, providing an evolutionary basis for virus-triggered vascular leak.

**Author Summary:** The *Flaviviridae* contain numerous medically important human pathogens that cause potentially life-threatening infections. Over half of the world’s population is at risk of flavivirus infection, and this number is expected to increase as climate change expands the habitats of the arthropod vectors that transmit these flaviviruses. There are few effective vaccines and no therapeutics approved for prevention or treatment of flavivirus infection, respectively. Given these challenges, understanding how and why flaviviruses cause pathogenesis is critical for identifying targets for therapeutic intervention. The secreted nonstructural protein 1 (NS1) of flaviviruses is a conserved virulence factor that triggers endothelial dysfunction in a tissue-specific manner. It is unknown if this endothelial dysfunction provides any benefit for virus infection. Here we provide evidence that NS1-triggered endothelial dysfunction facilitates virus crossing of endothelial barriers and augments infection of target cells *in vitro* and promotes virus dissemination *in vivo*. This study provides an evolutionary explanation for flaviviruses evolving the capacity to trigger barrier dysfunction and highlights NS1 and the pathways governing endothelial dysfunction, as therapeutic targets to prevent flavivirus dissemination.

## Introduction

The *Flaviviridae* family comprises many important human pathogens, including the four dengue virus serotypes (DENV1-4), Zika virus (ZIKV), West Nile virus (WNV), Japanese encephalitis virus (JEV), and yellow fever virus (YFV). More than half of the world’s population is at risk of flavivirus infection given the global distribution of the mosquito vectors that transmit these viruses.^1^ Flavivirus infections can be subclinical or display a range of clinical manifestations, from non-severe febrile illness to severe, potentially life-threatening disease associated with vascular leak.^1–3^ Severe DENV and YFV infection is associated with systemic vascular leak resulting in hypovolemic shock that can lead to multiorgan failure.^4–6^ WNV and JEV infections are neurotropic, and infection is associated with neurological pathologies including meningomyeloencephalitis.^1,2,7^ ZIKV causes congenital Zika syndrome in pregnant women resulting in microcephaly in infants, as well as being associated with Guillain-Barré syndrome in adults.^1,2,8^ Mechanistically, the triggers of these specific pathologies are unclear and are largely associated with an uncontrolled inflammatory response.^1,2^ Recently, the secreted flavivirus nonstructural protein 1 (NS1) has been shown to be a contributing factor to tissue-specific flavivirus pathogenesis.^9^

Flaviviruses are enveloped viruses with a positive-strand RNA genome of ~11 kb encoding for 3 structural proteins and seven nonstructural proteins, including NS1.^3^ NS1 is found within infected cells as a dimer associated with the replication complex and is critical for viral replication and egress.^10,11^ NS1 is also secreted from infected cells as higher-order oligomers (tetramers and hexamers) associated with lipid cargo and components of high-density lipoparticles.^12–16^ Levels of soluble NS1 circulating in the blood have been found to correlate with flavivirus disease severity.^17–19^ NS1 has been shown to function as a virulence factor in numerous ways, including by activating inflammatory responses, inhibiting complement, activating platelets, and inhibiting function of immune cell populations.^20–23^ Interactions between endothelial cells and flavivirus NS1 have also been shown to be a direct trigger of endothelial dysfunction and vascular leak.^9,24,25^ Mechanistically, the ability of NS1 to trigger vascular leak is dependent upon NS1 internalization into endothelial cells and subsequent disruption of key endothelial cell barrier components, including the endothelial glycocalyx (EGL) and intercellular junctions (IJC).^9,24,26,27^ Importantly, vaccination against NS1 or passive transfer of anti-NS1 antibodies have been shown to be protective against lethal flavivirus challenge.^20,28–33^

Flavivirus infection is initiated by the bite of an infected vector, starting in the skin and then moving to the blood before disseminating from the blood into specific tissues associated with disease. Vascular leak is a hallmark of severe dengue and yellow fever, and tissue-specific endothelial dysfunction in the brain and placenta are associated with WNV and ZIKV infection.^1,2,5,6^ While it is still unclear what factors influence this disease tropism, our recent work demonstrated that flavivirus NS1 triggers endothelial dysfunction and vascular leak in a tissue-specific manner that mirrors disease tropism of the respective flavivirus.^9,24,34^ For example, DENV NS1 triggers endothelial dysfunction on multiple tissue-derived human endothelial cell lines, while WNV and JEV NS1 trigger endothelial dysfunction only on brain-derived endothelial cells lines, correlating with systemic and neurotropic pathologies of DENV and WNV/JEV infections, respectively.^9,35^ This is mirrored *in vivo* in a wild-type mouse model by DENV NS1 induction of vascular leak in the lung, liver, and brain, while WNV NS1 only induces leak in the brain.^9^ Together, these data suggest that the ability of flavivirus NS1 to trigger tissue-specific endothelial dysfunction may influence virus tissue tropism. We hypothesize that the capacity of flavivirus NS1 to trigger vascular leak may promote virus infection of a host by mediating virus dissemination from the blood and into specific tissues. Indeed, there are studies supporting this hypothesis that demonstrate the role of NS1 in promoting dissemination of ZIKV from the blood into the testis as well as WNV from the blood into the brain.^36,37^

Here, we demonstrate the capacity of DENV NS1 to promote dissemination of DENV from the blood into the lungs of infected mice. *In vitro*, we monitor flavivirus dissemination across monolayers of endothelial cells and show that NS1 promotes this process. We also show that this process is complex, involving both a virus-independent diffusion across barriers and a virus-specific increase in infectivity of cells on the other side of endothelial barriers in an NS1-tissue-specific manner, suggesting a potential homotypic interaction between NS1 and flavivirus virions. We also provide evidence of NS1-virion interactions. Taken together, this study demonstrates that DENV NS1 promotes dissemination of DENV *in vivo* and creates a system to mechanistically investigate dissemination *in vitro*.

## Results

### DENV NS1 promotes DENV dissemination *in vivo*

We previously demonstrated the capacity of exogenous DENV NS1 to trigger vascular leak and the ability of passively transferred anti-NS1 antibodies to protect against vascular leak and morbidity/mortality in mouse models of DENV infection and disease.^28,29^ Despite these observations, it is unclear how NS1-triggered pathology modulates DENV infection *in vivo*. We hypothesized that the anti-NS1 monoclonal antibody (mAb) 2B7^29^ would block the capacity of DENV to disseminate into the lungs of mice and, conversely, that the addition of exogenous NS1 would promote dissemination of DENV into the lungs. We examined the lungs of mice because vascular leak in the lungs of humans, manifesting as pleural effusion, is a common feature of severe dengue.^9,28^ To test this hypothesis, we infected mice with a sublethal dose of a mouse-adapted strain of DENV2 (D220), co-administered the anti-NS1 protective antibody 2B7, then measured accumulation of DENV in the lungs of infected mice **(Fig. 1A)**. We found higher levels of DENV genome copies in mice that received an IgG isotype control compared to mice receiving a dose of 2B7, supporting our hypothesis **(Fig. 1A)**. Further, we found that the amounts of virus in the lungs were greater in mice receiving a dose of 100 µg of DENV NS1 at the time of virus infection compared to mice receiving a dose of 100 µg of ovalbumin **(Fig. 1B)**. Taken together, these data support the hypothesis that NS1-triggered vascular leak *in vivo* promotes dissemination of DENV into target tissues.

**Figure 1.**
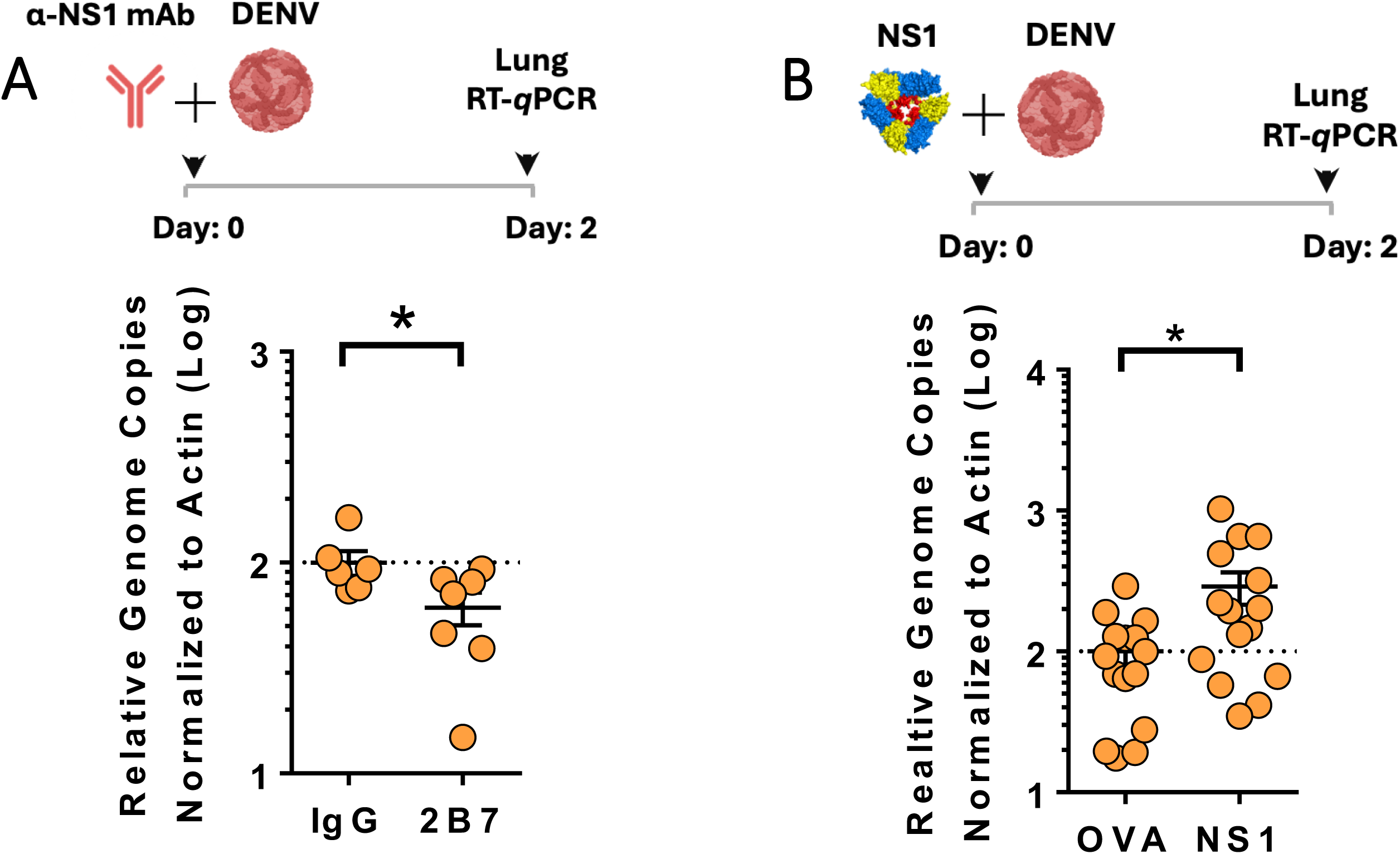
DENV NS1 promotes viral dissemination *in vivo*. **(A)** Viral dissemination assay measuring viral load in the lungs of C57BL/6 *Ifnar^−/−^* mice. Mice were administered 150 µg of the anti-NS1 antibody 2B7 or an IgG isotype control antibody via intraperitoneal injection and then infected with the mouse-adapted DENV2 strain D220 intravenously. Forty-eight hours post-infection, mice were sacrificed, lungs were collected, and viral RNA was quantified by RT-qPCR and normalized to levels of β-actin mRNA. **(B)** Same procedure as A, but mice were treated with 100 µg of DENV2 NS1 or ovalbumin as a negative control. Data are graphed as Relative Genome Copies Normalized to Actin with *p<0.05.

### NS1-triggered endothelial hyperpermeability promotes flavivirus dissemination across endothelial barriers

Our *in vivo* data suggest that NS1-triggered endothelial dysfunction promotes virus dissemination across endothelial barriers, but the nature of this dissemination is unclear. We hypothesized that virus passage across endothelial barriers would correlate with the extent of endothelial hyperpermeability triggered by NS1. To test this, we developed an *in vitro*-based dissemination assay to monitor the passage of virus across monolayers of endothelial cells seeded in the apical chamber of 24-well transwell inserts with semi-permeable membranes. Flavivirus NS1 (from DENV, ZIKV, WNV, JEV, and YFV) and distinct flaviviruses (DENV, ZIKV, and the West Nile strain Kunjin virus [KUNV]) were added to apically seeded endothelial cells derived from the lung (human pulmonary microvascular endothelial cells [HPMEC]), brain (human brain microvascular endothelial cells [HBMEC]), or umbilical vein (human umbilical vein endothelial cells [HUVEC]). We then monitored endothelial hyperpermeability using a transendothelial electrical resistance assay (TEER) and virus passage to the basolateral chamber using RT-qPCR **(Fig. 2A)**. As seen previously, we observed tissue-specific endothelial hyperpermeability induced by distinct flavivirus NS1 proteins. DENV NS1 and YFV NS1 (to a lesser extent) triggered endothelial hyperpermeability on HPMEC **(Fig. 2B)**.^9^ DENV and ZIKV NS1 triggered endothelial hyperpermeability on HUVEC **(Fig. 2C)**. DENV, ZIKV, WNV, and JEV NS1 triggered endothelial hyperpermeability on HBMEC monolayers **(Fig. 2D)**. Conducting RT-qPCR on media from the basolateral chamber at 24 hours after NS1-treatment and infection, we found that NS1 proteins that trigger endothelial hyperpermeability in the distinct endothelial cell lines promoted passage of virus across endothelial cell monolayers, correlating with our TEER data and supporting our hypothesis **(Fig. 2E-M)**. To confirm that the virus detected in the basolateral chamber was indeed input virus that had disseminated across the endothelial barrier, and not progeny virus resulting from infection of the endothelial cell monolayers, we monitored infection levels of each of the three endothelial cell lines by immunofluorescence microscopy. Infection of highly susceptible Vero-CCL81 cells served as a positive control. We found no evidence of viral antigen in HPMEC, HBMEC, or HUVEC for any virus infection 24 hours post-infection **(S1A Fig)**. In contrast, we detected high levels of viral antigen in Vero-CCL81 cells infected for 24 hours **(S1A Fig)**. Overall, these data indicate that flavivirus NS1 can promote passage of virions across cells in a virus non-specific manner.

**Figure 2.**
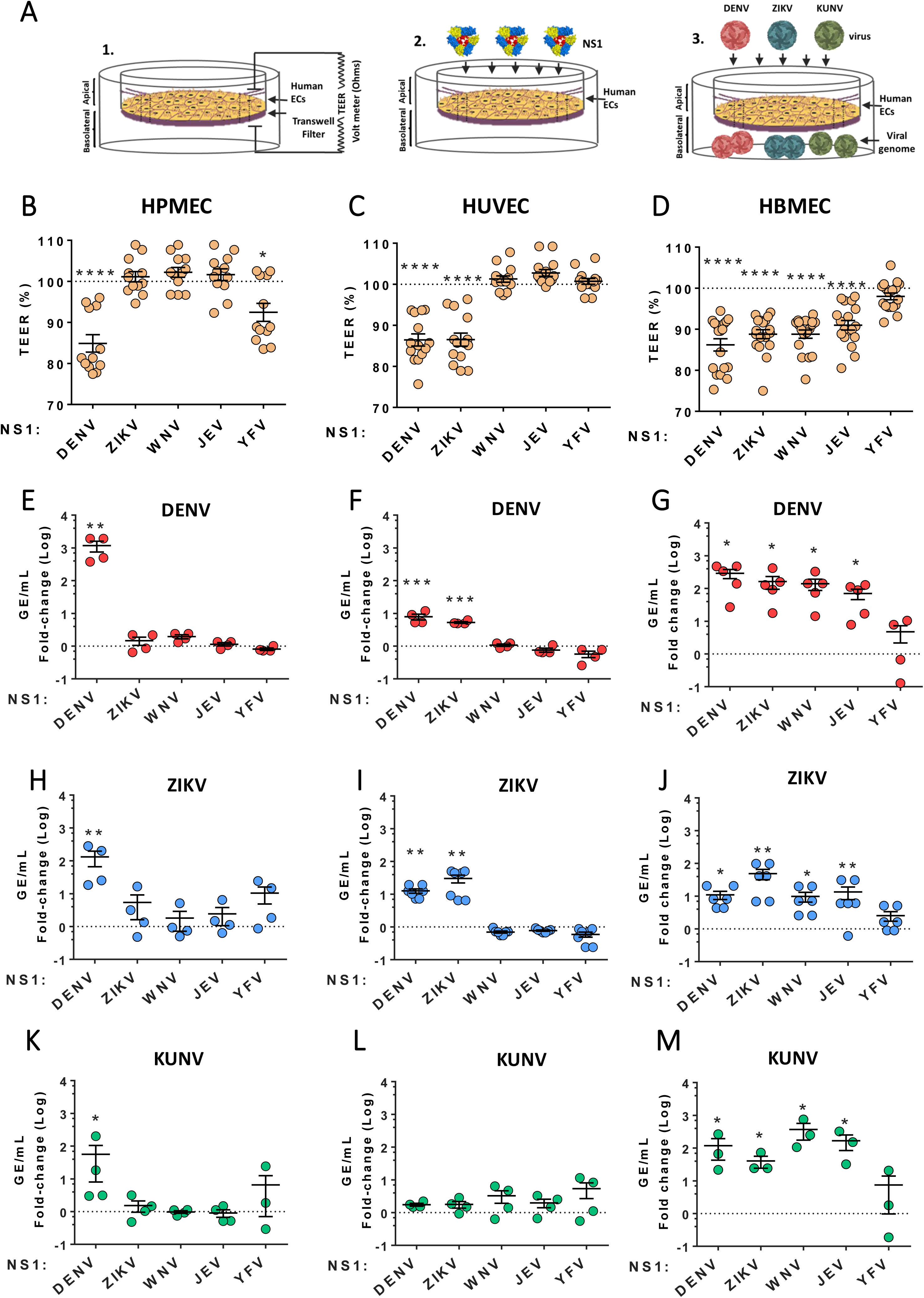
Flavivirus NS1 facilitates virus crossing of endothelial cell barriers. **(A)** Schematic of a Transwell-based system to measure (1 and 2) hyperpermeability and (3) virus crossing of endothelial cell monolayers seeded in the apical chambers of 24-well transwell inserts. **(B)** Transendothelial electrical resistance assay of HPMEC treated with 5 µg/mL of the indicated flavivirus NS1 protein, measured 3 hours-post treatment. The dotted line is the baseline of untreated cells. **(C)** Same as B, but with HUVEC. **(D)** Same as B, but with HBMEC. **(E-M)** The indicated viruses were added to the apical chamber of cells from B-D, and viral load was measured in the basolateral chamber 24 hours later by RT-qPCR. Panels E, H, and K are HPMEC treated with 1×10^5^ focus-forming units (FFU) of DENV, 1×10^5^ FFU of ZIKV, and 1×10^6^ FFU of KUNV, respectively. Panels F, I, and L received the same virus treatments, but with HUVEC. Panels G, J, and M received the same virus treatments, but with HBMEC. Data are expressed as fold-change (Log_10_) of genome equivalents (GE/mL) obtained with normalization to viral levels in the basolateral chamber of untreated (no NS1) endothelial cells. The mean +/− standard error of the mean are displayed for at least three biological replicates. One-way ANOVA with multiple comparisons statistical tests were conducted, with *p<0.05, **p<0.01, ***p<0.001, and ****p<0.0001.

We next modified our dissemination assay by seeding the monocytic cell line U937-DC-SIGN in the basolateral chamber of transwells and measured infection levels of these target cells by flow cytometry 24 hours after infection **(Fig. 3A)**. This allows us to measure not only viral particles crossing endothelial barriers, but also the capacity of these viruses to infect cells across endothelial barriers. As before, we hypothesized that the capacity to infect target U937-DC-SIGN would correlate with the level of NS1-triggered endothelial hyperpermeability. Surprisingly, while there was overall more infection of U937-DC-SIGN cells when endothelial cell barriers were compromised, we observed the greatest enhancement of infection when the virus was paired with its homotypic NS1 protein. For example, on HPMEC, while DENV NS1 promoted some infection of U937-DC-SIGN cells by ZIKV and KUNV, the most striking increase in infection was observed for DENV **(Fig. 3B, 3E, and 3H).** On HUVEC, DENV NS1 promoted the greatest target cell infection for DENV and ZIKV NS1 promoted the greatest infection enhancement for ZIKV, while KUNV infection of U937-DC-SIGN cells was only slightly enhanced in both DENV and ZIKV NS1 treatment conditions **(Fig. 3C, 3F, and 3I)**. For HBMEC, DENV NS1 most significantly modulated DENV infection, ZIKV NS1 enhanced ZIKV infection to the greatest extent, and WNV NS1 augmented KUNV infection to the greatest extent **(Fig. 3D, 3G, and 3J)**.

**Figure 3.**
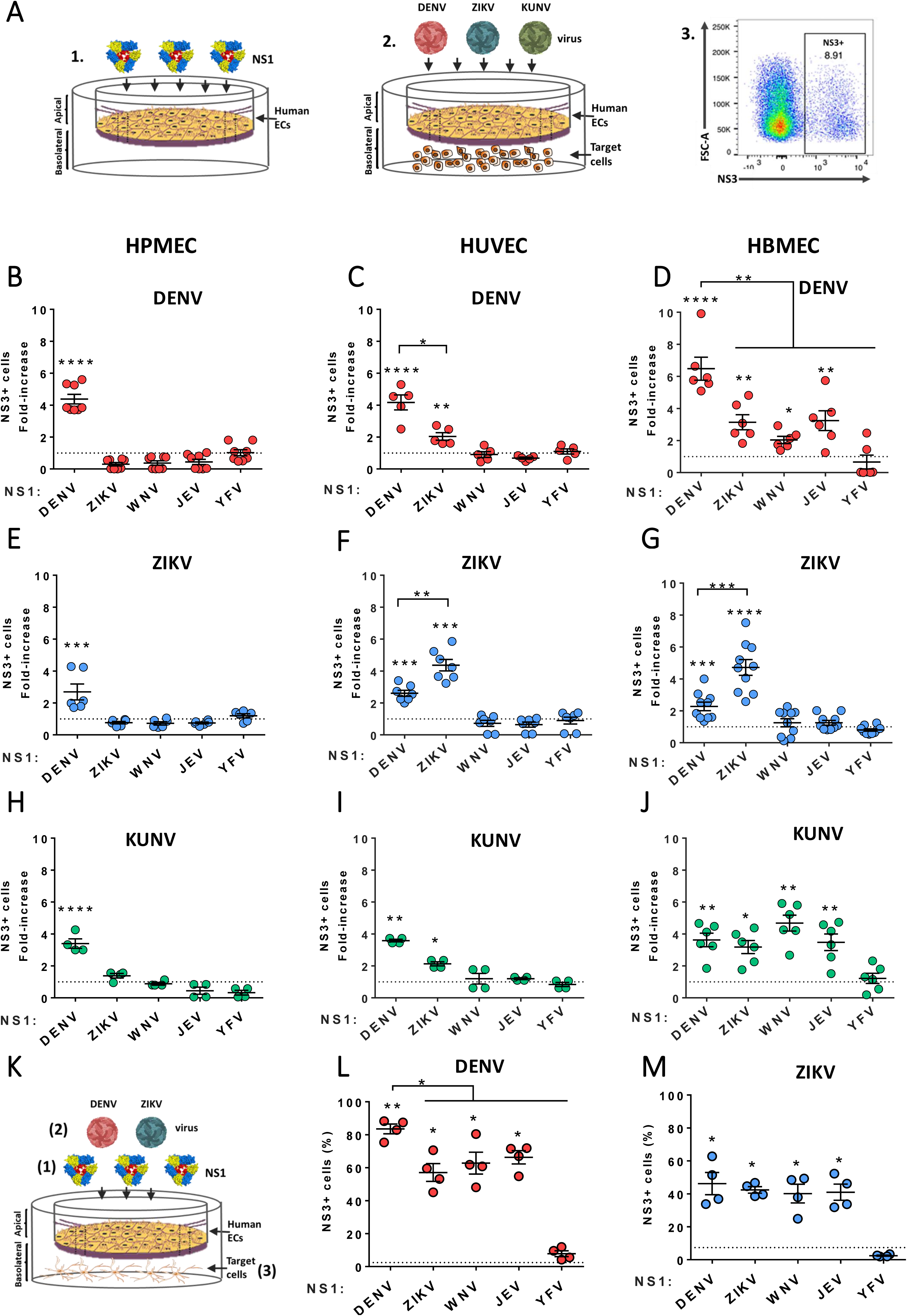
Flavivirus NS1 promotes infection of target cells across endothelial cell barriers. **(A)** Monolayers of human endothelial cells from lung (HPMEC, *left panel*), umbilical vein (HUVEC, *middle panel*), and brain (HBMEC, *right panel*) grown on transwells (1) were treated for 3 hours with different flavivirus NS1 proteins (5 µg/mL) (2). Target cells (human monocytic U937-DC-SIGN cells) were plated in the basolateral chamber of transwells (3). Then, a viral inoculum of purified **(B-D)** DENV (1 × 10^5^ FFUs), **(E-G)** ZIKV (1 × 10^5^ FFUs), or **(H-J)** KUNV (1 × 10^6^ FFUs) was added to the apical chamber. The percentage of infected cells on the basolateral side of transwells was determined by flow cytometry 24 hours later. Results are expressed as the fold-increase of target infected cells obtained between each experimental condition and the non-treated monolayers used as controls to measure background levels of virus crossing. Data represent the mean +/− error of the mean of at least n=3 biological replicates. **(K-M)** Normal human astrocytes (NHA) were plated in the basolateral chamber of HBMEC grown on transwells and infections were performed as above in the presence or absence of flavivirus NS1 proteins. The percentage of DENV- or ZIKV-infected NHA were examined by immunofluorescence staining for NS3 after 48 hours. Data represent mean +/− error of the mean of at least three biological replicates. One way ANOVA and non-parametric tests including Kruskal-Wallis and Mann-Whitney tests were performed for comparison between several groups and two groups, respectively. Statistically significant differences were considered as *p<0.05, **p<0.01, ***p<0.001, and ****p<0.0001.

In addition, we modeled the blood brain barrier by using HBMEC seeded in transwell apical chambers with human astrocytes seeded in the basolateral chamber **(Fig. 3K)**. Using immunofluorescent microscopy to visualize virus-infected astrocytes, we quantified infection following a similar protocol to the one described above. Comparably to what we observed before, while DENV infection was increased over background when HBMECs were treated with ZIKV, WNV, or JEV NS1, we found the greatest increase of astrocyte infection for DENV was when HBMEC layers were treated with the homotypic DENV NS1 **(Fig. 3L)**. For ZIKV infection, we found that NS1 from DENV, ZIKV, WNV, and KUNV all equally enhanced ZIKV infection, potentially because of high virus permissiveness of these human astrocytes to ZIKV infection **(Fig. 3M**).

### Antibodies targeting NS1 block dissemination of flaviviruses across endothelial barriers

To confirm the specificity of our NS1-triggered dissemination phenotype, we used antibodies targeting DENV or ZIKV NS1. We utilized the same transwell system with HUVEC seeded on the apical chamber and target U937-DC-SIGN cells seeded in the basolateral chamber. For this experiment, NS1 was precomplexed (or not) with the indicated mAbs before addition to the apical chamber **(Fig. 4A)**. For DENV NS1, we utilized two anti-NS1 mAbs with the capacity to block NS1 function (10E6 and 2B7) and a control mAb that does not block NS1-triggered endothelial hyperpermeability (2E9.E2) or an isotype control.^28,29^ As expected, we found that mAbs 10E6 and 2B7 abrogated the capacity of NS1 to trigger endothelial hyperpermeability of HPMEC **(Fig. 4B)**. Correlating with this anti-NS1 activity, we found that 2B7 and 10E6 blocked DENV infection of U937-DC-SIGN, comparable to cells not treated with NS1, and in contrast to cells treated with NS1 and given control antibodies or no antibody **(Fig. 4D)**. For ZIKV NS1, we used three mAbs (ZKA25, ZKA53, and ZKA39) in addition to an isotype control.^38^ We found that ZKA25 and ZKA53 abrogated the capacity of ZIKV NS1 to trigger endothelial hyperpermeability of HBMEC, in contrast to ZKA39 and the isotype control antibody **(Fig. 4C)**. Correlating with this blocking activity, we found that ZKA25 and ZKA53 blocked NS1-mediated ZIKV infection across monolayers of HBMEC, in contrast to ZKA39 and the isotype control antibody **(Fig. 4E)**. Taken together, these data indicate that NS1 specifically promotes increased infection of target cells by flaviviruses across monolayers of endothelial cells.

**Figure 4.**
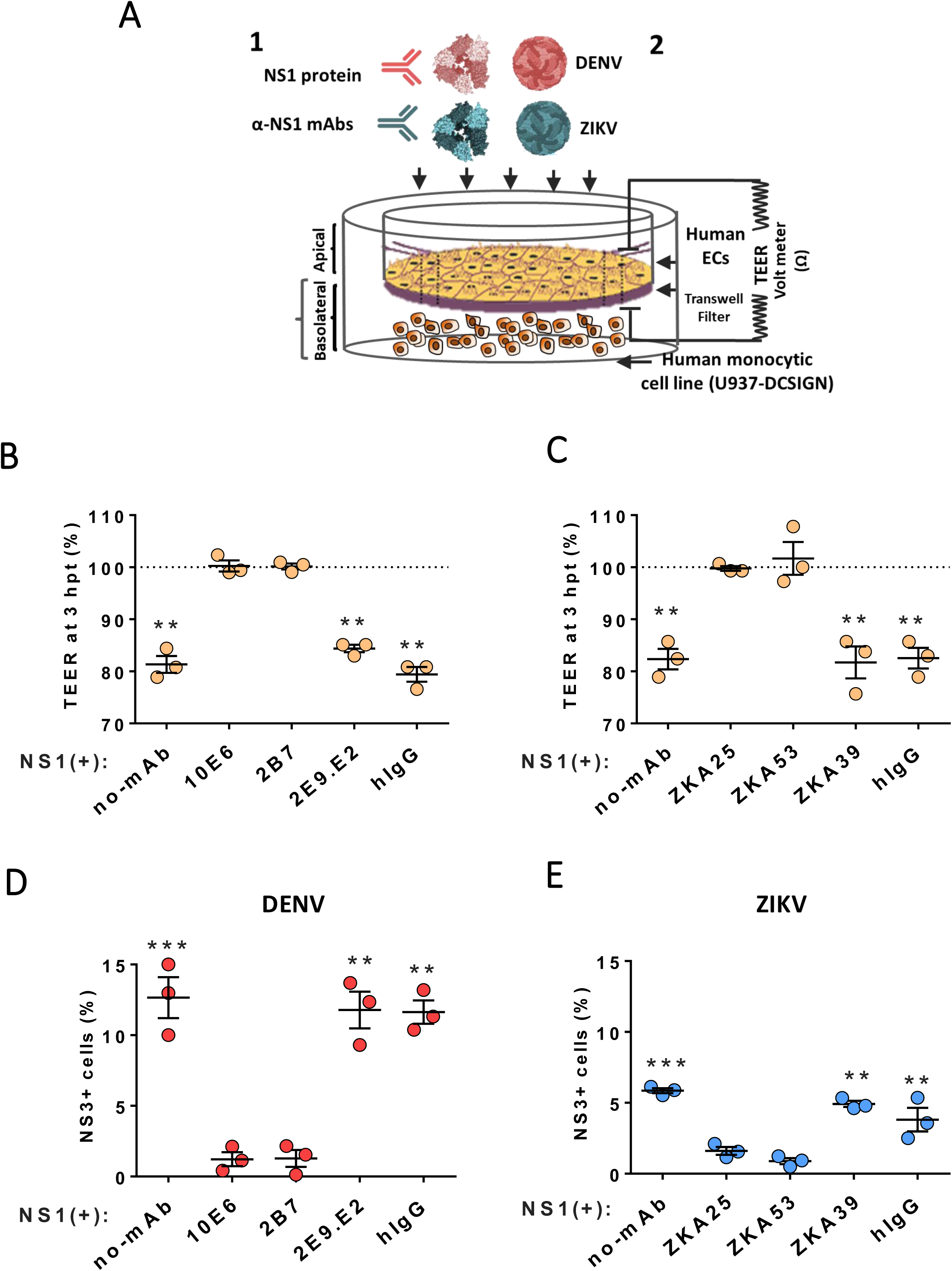
Anti-NS1 antibodies block NS1-induced endothelial hyperpermeability and virus dissemination across endothelial cell barriers. **(A)** Monolayers of HUVEC grown on transwells were treated for 3 hours with DENV and ZIKV NS1 proteins (5 µg/mL) in the presence and absence of 10 µg/mL anti-NS1 mAbs or human IgG as a negative control (1). Trans-endothelial electrical resistance (TEER) was used to measure the effect of anti-DENV NS1 **(B)** or anti-ZIKV NS1 **(C)** mAbs on DENV or ZIKV NS1-induced permeability of HUVEC monolayers, respectively. Then, purified DENV (1 × 10^5^ FFUs) or ZIKV (1 × 10^5^ FFUs) was added to the apical side of the monolayers **(A)** (2). The effect of anti-DENV NS1 **(D)** or ZIKV NS1 **(E)** mAbs on virus dissemination through HUVECs was examined by detecting the infection of target cells (U937-DC-SIGN cells) plated in the basolateral chamber of transwells. The percentage of infected cells on the basolateral side of transwells was determined by flow cytometry after 24 hours. Data represent the mean +/− standard error of the mean of the fold-change between NS1-treated and non-treated monolayers. Data are from n=3 biological replicates. One-way ANOVA and non-parametric tests including Kruskal-Wallis and Mann-Whitney tests were performed for comparison between several groups and two groups, respectively. Statistically significant differences were considered as **p<0.01 and ***p<0.001.

### NS1 associates with flavivirus virions

Our *in vitro* dissemination system indicates that NS1 both promotes passage of flavivirus virions across endothelial barriers and enhances infection of target cells. NS1 enhancement of infection was most pronounced when homotypic pairs of NS1 and flavivirus were used (*e.g.,* DENV NS1 and DENV virions). This suggests a potential interaction between virus particles and NS1. To determine if NS1 associates with flavivirus virions, we conducted a co-precipitation assay where 6xHIS-tagged NS1 proteins (from DENV or ZIKV) were preincubated with nickel resin and then mixed with DENV or ZIKV virions. Preincubation of nickel resin with BSA or virus alone served as negative controls. The resin was subsequently pelleted and washed, and co-precipitated virus was quantified by RT-qPCR **(Fig. 5A)**. We found that resin mixed with DENV and ZIKV NS1 contained more DENV virions compared to BSA and virus-only conditions **(Fig. 5B)**. Resin mixed with DENV NS1 and ZIKV NS1 co-precipitated higher levels of ZIKV virions, compared to other conditions **(Fig. 5C)**. Of note, we observed a significantly higher amount of virus precipitation when homotypic pairs of NS1 and virus were used **(Fig. 5B and 5C),** supporting the hypothesis of an interaction between flavivirus virions and homotypic NS1 proteins.

**Figure 5.**
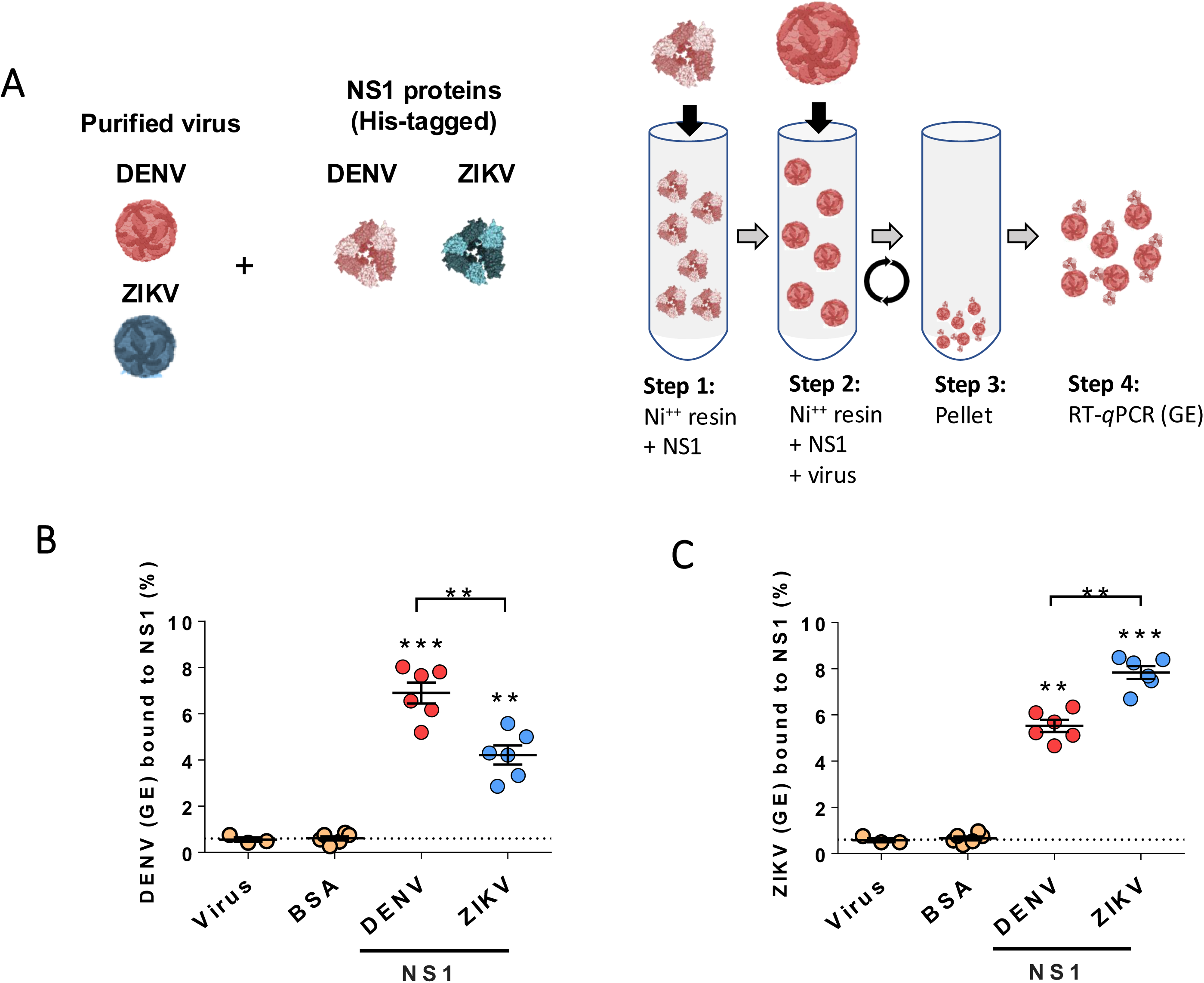
DENV and ZIKV particles co-precipitate with NS1 proteins. **(A)** DENV and ZIKV NS1 (containing C-terminal 6xHIS tags) were diluted to 5 μg in NTE buffer, then mixed with Ni-NTA resin for 2 hours at 4°C rotating (*Step 1*). NS1-bound resin was washed 2x with 1X PBS, then purified virus (DENV or ZIKV, 1 × 10^9^ viral particles) was added to the resin and allowed to bind for 2 hours at 4°C rotating (*Step 2*). Beads were washed 2x with cold NTE buffer, then 2x with 30 mM Imidazole in NTE buffer, then eluted with 300 mM Imidazole (*Step 3*). Eluted NS1 and virus was collected, and viral RNA (Genome Equivalents, GE) was purified and quantified by RT-qPCR (*Step 4*). **(B)** RT-qPCR detecting DENV RNA (GE) from eluates of Ni-NTA resin bound to the indicated proteins. BSA and virus alone conditions serve as negative controls for background levels of virus binding. Results are displayed as percent of virus bound from total input virus (GE). Data are from at least n=3 biological replicates. One-way ANOVA and non-parametric tests including Kruskal-Wallis and Mann-Whitney tests were performed for comparison between several groups and two groups, respectively. Statistically significant differences were considered as **p<0.01, and p***p<0.001.

We hypothesized that the dual activities of NS1 (*i.e.,* promoting dissemination across barriers and modulating virus infectivity) would utilize distinct regions of NS1. Flavivirus NS1 contains three domains: the β-roll, wing, and β-ladder.^35,39^ We have previously demonstrated that the β-ladder is critical for DENV NS1 to trigger endothelial dysfunction and vascular leak, with numerous key amino acid residues identified, including N207.^26,29,35^ Further, our biochemical investigation of the association of flavivirus virions and NS1 identified DENV NS1 residue 343 within the β-ladder as a potential meditator of virion association.^40^ Here, we tested the relative contribution of residues N207 and E343 for NS1 to facilitate virus dissemination. Using the same experimental design described above, we found that the DENV NS1-E343K mutant protein displayed a diminished capacity to precipitate DENV virions compared to DENV NS1-WT, but greater than BSA and virus alone control conditions **(Fig. 6A)**. This experiment suggested that NS1-E343K possesses a defect in the capacity to associate with virions. To determine the mechanisms of this defect, we then compared the capacity of NS1-WT and NS1-E343K to interact with endothelial cells and trigger endothelial hyperpermeability of HPMEC. We found that NS1-E343K bound to HPMEC at levels comparable to NS1-WT, suggesting that the defect of NS1-E343K is not explained by a defect in endothelial cell interaction **(Fig. 6B)**. We next conducted TEER on HPMEC treated with NS1-WT, NS1-N207Q, and NS1-E343K. As we previously demonstrated, NS1-WT triggered endothelial hyperpermeability, in contrast to NS1-N207Q, which did not alter barrier function.^26^ Interestingly, we found that NS1-E343K could still trigger endothelial hyperpermeability at levels comparable to NS1-WT **(Fig. 6C)**, suggesting that the capacity to interact with virions and trigger endothelial hyperpermeability may be distinct properties mediated by different regions of NS1. Despite this differential activity, we found that both NS1-N207Q and NS1-E343K were unable to enhance infection of U937-DCSIGN target cells across barriers at the same rate as NS1-WT, suggesting that both the capacity of NS1 to trigger endothelial hyperpermeability and to interact with virions are required for dissemination **(Fig. 6D)**.

**Figure 6.**
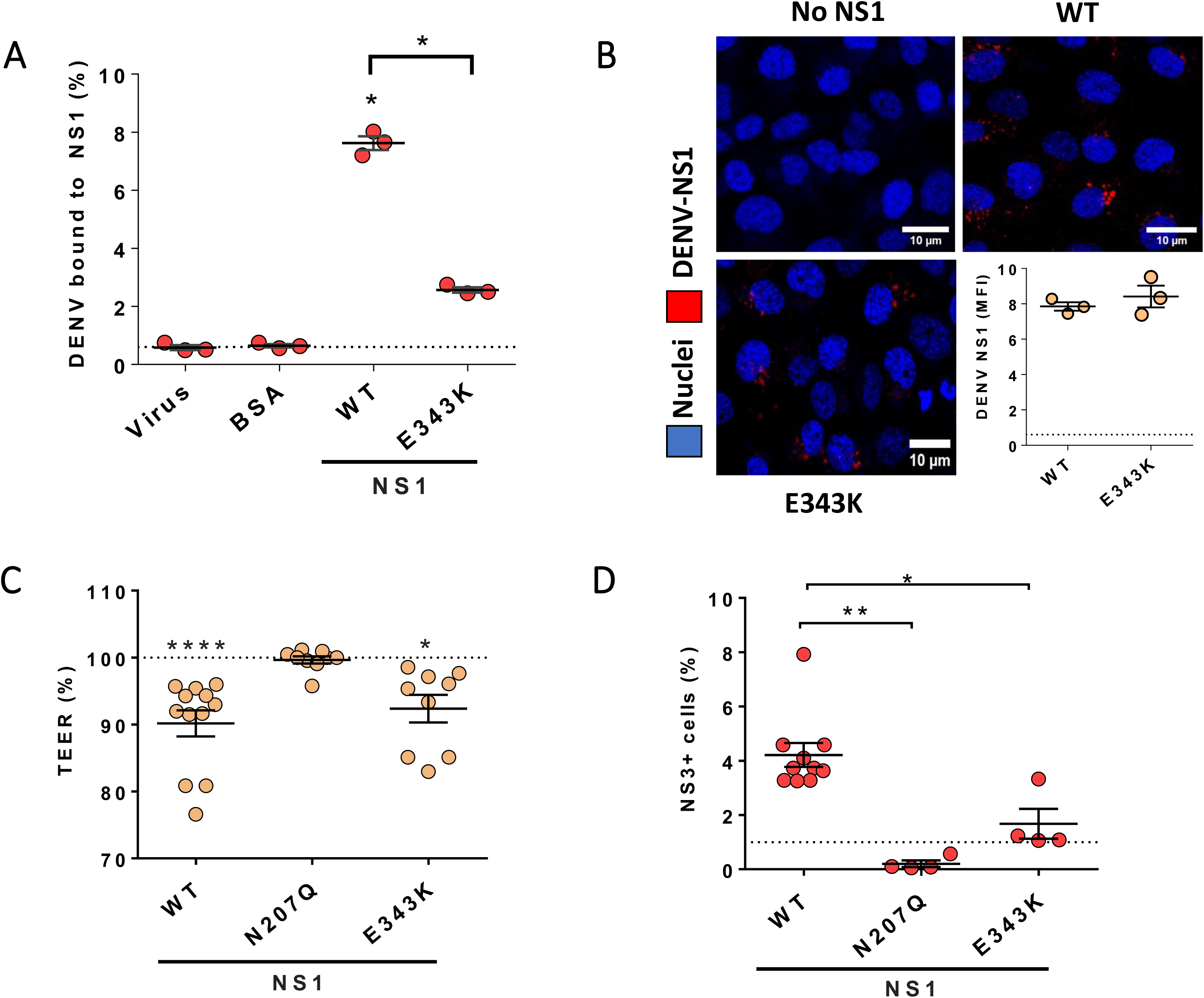
DENV NS1-E343K does not co-precipitate with DENV virions or mediate enhanced infection of target cells *in vitro*. **(A)** NS1-WT or NS1-E343K (5 μg/mL) were precipitated with Ni-NTA resin for 2 hours then incubated with 1×10^9^ purified DENV genome equivalents for 2 hours at 4°C. The resin was then washed, and bound NS1 was eluted. Bovine serum albumin (BSA) or virus alone served as negative controls. Levels of eluted virus were quantified by RT-qPCR. **(B)** Binding of the indicated in-house-produced NS1 proteins (10 μg/mL) to HPMEC one hour post treatment analyzed by immunofluorescent microscopy assay. Data are mean fluorescence intensity of NS1 signal analyzed using ImageJ software. Red is NS1 (6xHIS) and blue is nuclei (Hoechst). **(C)** A transendothelial electrical resistance assay of HPMEC treated with the indicated in-house-produced DENV NS1 proteins (5 μg/mL) at 3 hours post-treatment. **(D)** The same cells as C were then treated with 1×10^5^ FFU of purified DENV added to the apical side. Twenty-four hours later, the percentage of infected cells (plated in the basolateral chamber) were determined by flow cytometry. In all figures, data represent the mean +/− standard error of the mean with at least three biological replicates. One-way ANOVA and non-parametric tests including Kruskal-Wallis and Mann-Whitney tests were performed for comparison between several groups and two groups, respectively, with *p<0.05; **p<0.01, and ****p<0.0001.

## Discussion

In this study, we investigated the role of NS1-mediated endothelial dysfunction in promoting flavivirus infection. A critical but complicated ongoing question is understanding why viruses cause disease. Is virus-triggered pathogenesis simply an “accident” or are pathogenic host pathways beneficial to virus infection? In the case of flavivirus infection, especially for DENV, vascular leak is a pathogenic hallmark of infection.^1,4,17^ Is this vascular leak simply an unintended consequence of viral infection or does vascular leak somehow benefit the virus? Here we find that the capacity of NS1 to trigger endothelial dysfunction and vascular leak promote dissemination of DENV into lungs of mice and dissemination of DENV, ZIKV, and KUNV across endothelial monolayers *in vitro*. This is consistent with other work demonstrating that WNV and ZIKV NS1 also promote virus dissemination into the brain and testis, respectively.^36,37^ Interestingly, NS1 also possesses the capacity to increase viral infection of target cells across endothelial barriers. Our study provides evidence that this increase in infectious capacity may be due to an association between homotypic pairs of virions and NS1 **(Fig. 7)**. Thus, our study provides an evolutionary explanation for the capacity of flaviviruses to trigger vascular leak. Although NS1 acts as a virulence factor that enhances pathogenesis in infected people, its role in promoting dissemination of virus across endothelial barriers may place evolutionary pressure on flaviviruses to preserve this function of NS1, as it presumably leads to higher viral load and higher viremia, which would increase transmission to mosquitoes.

**Figure 7.**
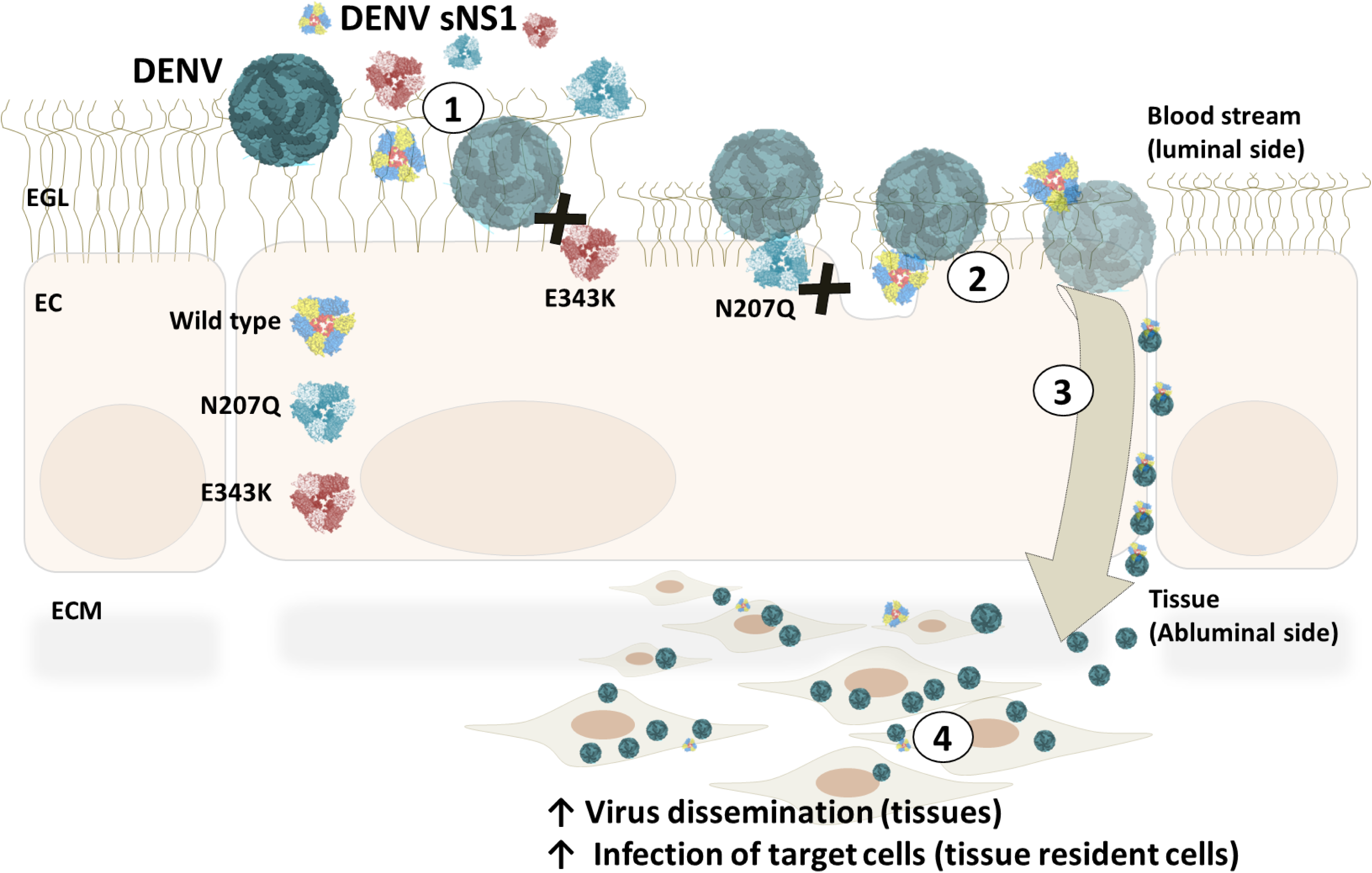
Flavivirus NS1 induces tissue-specific endothelial hyperpermeability facilitating viral dissemination and promoting infection. Flavivirus NS1 interacts with endothelial cells resulting in endothelial barrier disruption (1). Our work suggests an association between flavivirus NS1 and flavivirus virions involving NS1 residue 343 (2). Collectively, our model proposes that NS1 facilitates flavivirus crossing of endothelial barriers (3) and enhances infection of target cells (4). We find that NS1 mutants with decreased capacity to trigger endothelial hyperpermeability (DENV NS1-N207Q) and decreased capacity to interact with virions (DENV NS1-E343K) display decreased capacity to mediate virus crossing and enhanced infection of target cells. These findings suggest a new mechanism for secreted viral proteins to facilitate virus dissemination into tissues to encounter target cells that support viral replication.

Flaviviruses face the common challenge of traversing endothelial barriers in infected humans when disseminating from the initial site of infection in the dermis into the blood (following an insect bite), then from the blood into distal tissues, resulting in high levels of viral replication, and then from the tissue and back into the blood to be taken up by mosquito to be transmitted. Our work supports the hypothesis that NS1-triggered endothelial dysfunction promotes the dissemination of virus from the blood into tissues, then presumably back into the blood after amplification. Previous work provides evidence that high NS1 levels in the blood can promote infection of mosquitoes following a blood-meal.^41,42^ The mechanisms by which NS1 promotes mosquito infection by flaviviruses is unclear, but it is tempting to speculate that NS1 may also modulate barrier function of cells within mosquitoes. Our study focuses on the role of NS1 in promoting dissemination from the blood into distal tissues in mice (the lung in the case of our experiments), since we delivered DENV intravenously. Our *in vitro* investigation provides mechanistic insight regarding NS1-facilitated virus dissemination, revealing dual pathways of enhancing virus crossing of barriers and modulation of infectivity of target cells across barriers **(Fig. 7)**. Future work will be required to examine the role of NS1 in promoting dissemination of dermally administered virus into the blood and test whether NS1 impacts barrier function of mosquito cells. Future studies will also be needed to probe how NS1 enhances infection of DENV in target cells and if this indeed occurs *in vivo*.

Our model predicts that the proviral role of extracellular NS1 may occur after a host is inoculated with a flavivirus following an insect bite. Specifically, while NS1-triggered barrier dysfunction has been associated with severe disease manifestations in animal models of infection and people, the tissue-specific endothelial dysfunction triggered by NS1 allows the virus to disseminate across barriers, gain access to virus-permissive cells, and promote infection of these cells, presumably to increase viral transmission, before severe disease manifestations occur. We speculate that the severe disease manifestations associated with NS1 are either an indirect result of the increased virus titers resulting from increased dissemination and/or the result of excess NS1 protein produced late in infection. It is important to consider that severe dengue is certainly a multifactorial process, and the relative contribution of NS1 to disease manifestations is not completely clear. Interestingly, our data indicates that antibodies targeting NS1 can block flavivirus dissemination. Thus, antagonizing NS1 not only serves to block NS1-triggered vascular leak late in infection, but also virus dissemination across barriers, further highlighting the promise of development of therapeutics targeting NS1 or including NS1 in flavivirus vaccination formulations.

Our study suggests that the capacity of NS1 to promote virus infection may be due to an association between NS1 and flavivirus virions. Our previous data indicate that residue 343 within the β-ladder of NS1 plays a role in this interaction^40^, and we can indeed show that the NS1-E343K mutant protein displays a decreased capacity to associate with virions or enhance infection of target cells across endothelial cell monolayers *in vitro*. Intriguingly, this mutant NS1 protein can still trigger endothelial dysfunction, suggesting that the capacity of NS1 to trigger endothelial dysfunction is not completely sufficient to promote viral dissemination. Further studies are warranted, including structural studies aimed at visualizing the putative interaction between flavivirus virions and their homotypic NS1 proteins. Interestingly, while the 343 residue is a part of the β-ladder domain of NS1, it is found on the polar face of NS1, in contrast to previous β-ladder mutations that we identified to be critical for triggering endothelial dysfunction that are located on the cell-facing surface of NS1.^29,35^ This would imply a model where regions of NS1 in proximity to endothelial cells contribute to NS1-triggered endothelial dysfunction, while residues exposed to the aqueous environment may be free for interactions with virions. Thus, NS1 may serve as a molecular bridge to aid virus passage across barriers and modulate infectivity **(Fig. 7)**.

Taken together, our study provides evidence that flavivirus NS1 promotes virus dissemination. Previous work provides *in vivo* evidence that WNV NS1 promotes dissemination of virus into the brain and that ZIKV NS1 promotes virus dissemination across the blood-testis barrier.^36,37^ Our study demonstrates that DENV NS1 promotes dissemination of the virus into the lung and provides mechanistic detail on this dissemination phenotype. Taken together, evidence is growing that flavivirus NS1 promotes tissue-specific passage of flaviviruses into target tissues, acting as a “viral toxin”. This implies that viral and disease tropism may be partially dictated by the activity of NS1. This may also be the case for viruses outside of the *Flaviviridae* family. For instance, our group has described SARS-CoV-2 Spike and Crimean-Congo hemorrhagic fever virus (CCHFV) GP38 as viral toxins that trigger vascular leak and, in the case of GP38, promote virus dissemination *in vivo*.^43,44^ While more work is needed to expand the new field of viral toxins, our study provides evidence that flavivirus NS1 both triggers vascular leak and promotes virus dissemination, making it an attractive target for future therapeutic and vaccination campaigns.

## Materials and Methods

### Cell lines and viruses

Human Pulmonary Microvascular Endothelial Cells (HPMEC) were a gift from Dr. J.C. Kirkpatrick at Johannes Gutenberg Germany. Human Brain Microvascular Endothelial Cells (HBMEC) were a gift from Dr. Ana Rodriguez at New York University and originally obtained from ScienCell Research Laboratories. Human Umbilical Vein Endothelial Cells (HUVEC) were a gift from Dr. Melissa Lodoen at the University of California, Irvine. All endothelial cells were grown in Endothelial Growth Medium-2 BulletKit^TM^ medium (EGM2) from Lonza and maintained at low passages as previously described.^9^ Normal human astrocytes (NHA) were commercially obtained (CC-2565, Lonza) and maintained using Astrocyte Growth Medium (AGM, Lonza) according to manufacturer’s instructions. The U937 human monocytic cell line expressing the flavivirus attachment factor, DC-SIGN (U937-DC-SIGN, hereafter referred to as U937) was cultured and maintained using RPMI 1640 medium (Gibco) supplemented with fetal bovine serum (FBS, 2%), penicillin/streptomycin (1%), and glutamine (1%). FreeStyle 293F suspension cells (obtained from Thermo Fisher Scientific) were used to produce recombinant NS1 proteins and were maintained in FreeStyle 293 Expression medium (Thermo Fisher Scientific) at 37°C and 8% CO2 at a density of 0.15×10^6^ − 1.2×10^6^ cells/mL on a cell shaker at 135 rpm. The *Aedes albopictus* mosquito cell line (C6/36) was obtained from the American Type Culture Collection (ATCC) and maintained at 30-32°C in 5% C0_2_ in DMEM or RPMI medium supplemented with 1% Pen-Strep and fetal bovine serum. C6/36 cells were used to propagate all virus stocks in this study. Vero green monkey kidney cells (Vero-CCL81 from ATCC) were maintained in DMEM supplemented with high glucose (4.5 g/L), L-glutamine, 2% FBS and 1% P/S. Mouse-adapted DENV2 (D220) was previously produced as described previously.^29^ DENV serotype 2 (N172-06) and ZIKV (Nica 2-16) strains were isolated by the National Virology Laboratory, Ministry of Health, Managua, Nicaragua, and propagated (low passage: 2–3) in C6/36 cells at the University of California (UC), Berkeley. Kunjin virus (KUNV MRM61C infectious clone)^45^, a genetically stable Australian flavivirus closely related antigenically to West Nile virus (WNV), was kindly donated by Richard Kuhn (Purdue University, IN) and propagated in C6/36 cells at UC Berkeley. All viral stocks were tested and confirmed to be *Mycoplasma*-free, and viral titers were determined by standard focus-forming assays on Vero-CCL81 cells.^46^ Each virus was purified following standard procedures (e.g., Optiprep) to remove NS1 from the supernantant.^47^ Briefly, cellular supernatant obtained from virus-infected C6/36 cells (75 cm^2^ tissue culture flask, Corning) with each virus (DENV, ZIKV, KUNV) was collected 5–8 days postinfection and concentrated using 100-kDa Amicon filter units (Millipore) by centrifuging at 3250xg for 20 min at 4°C. The concentrated supernatant containing the virus was layered on a discontinuous 20/55% OptiPrep (Sigma-Aldrich) density gradient and ultracentrifuged at 40,000 rpm for 2 hours at 4°C, without brake, using a SW41Ti rotor. DENV layered between the 20 and 55% gradients was collected and tested by RT-PCR.

### Recombinant proteins and antibodies

NS1 proteins from distinct flaviviruses including DENV (DENV2 Thailand/16681/84), Zika virus (ZIKV Suriname Z1106033), West Nile virus (WNV NY99), Japanese Encephalitis virus (JEV SA-14), and Yellow Fever virus (YFV 17D) were obtained from The Native Antigen Inc. (Oxford, England), composed mostly of oligomeric forms with a purity greater than 95%, and certified by the manufacturer to be free of endotoxin contaminants.^29^ Imject^TM^ Ovalbumin (ThermoFisher Scientific) and an immunoglobulin G isotype control (ChromPure™ Human IgG, whole molecule, Jackson ImmunoResearch, Laboratories Inc.) were purchased and used as negative controls throughout this study. In-house produced WT and mutant DENV NS1 proteins, including NS1-N207Q and NS1-E343K, were produced by transfecting 293F cells with pMAB plasmids expressing the 6x-HIS tagged NS1 protein of interest. NS1 was purified from supernatants using cobalt resin and eluted using imidazole-containing elution buffer. Purified NS1 proteins underwent buffer exchange into 1x PBS and were determined to be highly pure and oligomeric using silver staining, Native-PAGE, and size exclusion-chromatography following standard procedures as previously described.^26,43^ One monoclonal antibody (mAb) was used to study flavivirus infection by immunofluorescence assay (IFA): the pan-flavivirus anti-non-structural protein 3 (NS3) antibody (E1D8).^48^ A goat anti-mouse IgG secondary antibody conjugated to Alexa 647 was used to detect viral infection (Thermo Fisher Scientific). For the NS1-blocking experiments *in vitro* and *in vivo* we used a panel of anti-NS1 mouse mAbs either generated in the Harris laboratory (P.R.B. and E.H.) including 2B7, 10E6 and 2E9.E2 or kindly donated from Davide Corti (Humabs BioMed) including the anti-ZIKV NS1 human IgG1 mAbs ZKA29, ZKA33, ZKA59 as previously published.^38^ For mouse mAbs, the hybridomas generated were propagated in CELLine culture flasks (Sigma); supernatants were collected, and the mAbs were affinity-purified using a Protein G Sepharose column. The mAbs were eluted from the column, concentrated, sterile-filtered, and titered before use.

### *In vivo* murine virus dissemination assay

All mice used in this study were bred in-house in compliance with Federal and University regulations. All animal experiments were approved by the Animal Care and Use Committee (ACUC) of UC Berkeley. For all experiments, six- to eight-week-old C57BL/6 Ifnar^−/−^ mice of both genders, housed under specific pathogen-free conditions, were transferred from the breeding facility to the infection facility. Mice were administered a dose of 150 μg of antibody or 100 μg of DENV NS1 (The Native Antigen Company) or Ovalbumin (OVA) (ThermoFisher Scientific) as indicated in the figures, via intravenous injection (IV) (NS1 and OVA) or intraperitoneal (IP) injection (antibodies). The next day, mice were infected IV with 1×10^5^ PFU of the DENV2 mouse-adapted strain D220. Forty-eight hours later (Day 2), mice were euthanized, and lung tissues were collected for virus quantification via RT-qPCR. Briefly, viral RNA was extracted from whole lungs using TRIzol (Invitrogen) according to the manufacturer’s specifications, and viral RNA levels were quantified using RT-qPCR. In brief, cDNA was reverse-transcribed from 1 µg of total RNA using IMPROM-II reverse transcriptase (Promega, A3803) with a random hexamer primer according to the manufacturer’s instructions. qPCR was conducted using SYBR-green reagents on an QuantStudio Real-Time PCR system. Relative genome copies of each condition were calculated through normalization to a house-keeping transcript encoding for beta-actin. The following primers were used for this study: D2_F_q_9944, 5’-ACAAGTCGAACAACCTGGTCCAT-3’; D2_R_q_10121, 5’-GCCGCACCATTGGTCTTCTC-3’; Actin-F, 5’-GCCTTCCTTCTTGGGTATGG-3’; Actin-R, 3’-GCACTGTGTTGGCATAGAGG-3’.

### Trans-endothelial electrical resistance (TEER) assay

TEER was used to investigate the impact of flavivirus NS1 proteins on the permeability of endothelial cells. In brief, endothelial cells were grown on transwell inserts in a 24-well Transwell polycarbonate membrane system (Transwell® permeable support, 0.4 μM, 6.5 mm insert; Corning Inc.) as previously described.^9^ Three hours post-treatment with 5 µg/mL of individual NS1 proteins (DENV, ZIKV, WNV, JEV, and YFV), TEER values were recorded as Ohms (Ω) using a manual electrode (EVOM2, World Precision Instruments) to measure the electrical resistance across monolayers of HPMEC, HUVEC, and HBMEC. Resistance values of transwells with no cells (blank) and untreated cells were used to calculate relative TEER as a ratio of the corrected resistance values as (Ω experimental condition - Ω blank)/(Ω untreated - Ω blank).

### Flow cytometry

The percentage of DENV-, ZIKV- or KUNV-infected suspension target cells (U937) plated in the basolateral side of transwell inserts containing monolayers of human endothelial cells was determined by flow cytometry using an anti-NS3 mAb (E1D8 conjugated to Alexa 647) to detect flavivirus-infected cells. Briefly, 24 hours post-infection, cell pellets were collected and washed 3X with pre-chilled FACS buffer (1X PBS, 2 mM EDTA, and 1% FBS) by centrifugation at 1,500 rpm at 4°C for 5 minutes. After fixation using 4% formaldehyde diluted in FACS buffer for 30 minutes at 4°C), cells were blocked and permeabilized using FACS buffer supplemented with 0.2% of saponin, 2% BSA and human whole IgG (10 μg/mL, Jackson ImmunoResearch) for 20 minutes at room temperature. In the same buffer, an optimized dilution of anti-NS3 mAb (E1D8 at 1:500) in-housed conjugated to Alexa 647 (Thermo Scientific) was added to each well and incubated with gentle rocking oscillation for 45 minutes, protected from light. Then cells were washed 2x using blocking/permeabilization buffer and then 2X with FACS buffer alone. After this, cells were acquired using a LSRFortessa multicolor flow cytometry at the UC Berkeley Core facility. Side scatter/Forward scatter (SSC/FSC) and singlet (SSC-A/SSC-H) parameters were used to gate the homogenous cell population visualized and normalized using counter density plots. Uninfected cells were used to determine the background from the staining protocol and to gate NS3+-positive cells as an indication of virus infection. Data analyses was performed using FlowJo software.

### Real-time RT-PCR

Detection and quantification of flavivirus RNA genomes in *in vitro* assays was performed by real-time RT-PCR (RT-qPCR). Briefly, 24 hours post-treatment with purified DENV, ZIKV, and KUNV, viral RNA was extracted using the QIAamp Viral RNA Mini Kit (Qiagen) from cell-free supernatants collected from the basolateral side of human endothelial cells grown on transwell inserts. Viral RNA was detected using the Verso 1-Step qRT-PCR kit (Thermo Fisher) in combination with distinct sets of primers and probes from previous studies). An NS5-targeted RT-qPCR assay for DENV was performed according to previous publication, with slight modification of the probe to recognize DENV2 N172-06 used in this study (DENV-F: 5′-F-ACAAGTCGAACAACCTGGTCCAT-3′; DENV-R: 5′-GCCGCACCATTGGTCTTCTC-3′; DENV-Probe: FAM-TGGGATTTCCTCCCATGATTCCACTGG-TAMRA). ZIKV primers and probe were modified from Lanciotti et al.,^49^ to recognize both Ugandan and Nicaraguan strains of ZIKV (ZIKV-F: 5′-F-CCGCTGCCCAACACAAG-3′; ZIKV-R: 5′-R-CCACTAACGTTCTTTTGCAGACAT-3′; ZIKV-Probe: FAM-AGCCTACCTTGACAAGCAATCAGACACTCAA-TAMRA). KUNV primers and probe were modified from previous publications to better recognize the KUNV strain used in this study (KUN-F: 5′-GGGCCTTCTGGTCGTGTTC-3′; KUN-R-5′-GATCTTGGCtGTCCACCTC-3′, KUNV-Probe: FAM-CCACCCAGGAGGTCCTTCGCAA-TAMRA). RNA standards generated by *in vitro* RNA transcription using the RiboMax kit (Promega P1300) were used to quantify viral RNA genomic equivalents at a range of 10^2^ to 10^8^ viral copies. DENV, ZIKV, and KUNV PCR amplicons were modified to include a T7 promoter by PCR, followed by *in vitro* transcription. Final RT-qPCR curve threshold (Ct) values were compared to a standard curve as described above, and total viral copy numbers are indicated as genomic equivalents (GE/mL). RT-qPCR assays were performed in a total volume of 30 µL containing 2X reaction mix (15µL), forward primer (10µM, 3µL), reverse primer (10µM, 3µL), enzyme mix (0.6 µL), nuclease-free water (4.5 µL), plus RNA template.

### Virus dissemination assay *in vitro*

To study the effect of NS1-induced endothelial hyperpermeability on the dissemination of DENV, ZIKV, and KUNV across monolayers of HPMEC, HUVEC, and HBMEC, two approaches were used. The first approach utilizes flow cytometry and immunofluorescence (IFA) assays to determine the percentage of DENV, ZIKV or KUNV infection of target cells (U937 or normal human astrocytes [NHA]) seeded in the basolateral side (bottom chamber) of endothelial cell monolayers grown on transwell inserts. The second approach uses real-time RT-qPCR to quantify the viral RNA GE for each flavivirus in the basolateral side of endothelial cells cultured on transwells. Regarding the first approach, 3 hours post-treatment with distinct NS1 proteins, and right before measuring TEER and adding the purified virus to the apical side of transwells, 500 μL of cell culture medium was removed from the bottom chamber of each transwell and replaced with 500 μL of fresh RPMI medium containing U937 cells (8×10^4^ cells) grown in suspension. In the case of NHA, transwells containing endothelial cells were carefully transferred on top of 24-well plates containing NHA (1×10^5^ cells) previously grown on (gelatin-treated (0.2%) coverslips until reaching 70-80% confluency using Astrocyte Growth Medium (AGM, Lonza). The percentage of infected target cells was determined by flow cytometry using E1D8 (anti-NS3 antibody) (see flow cytometry assay). Staining of the NS3 viral protein was performed in NHA 48 hours post-infection using previously described protocols for IFA staining.^25^ The percentage of infection with DENV and ZIKV was determined after counting the number of NS3 positive cells and the total number of cells per quadrant taken as 100%.

Quantification of viral particles crossing endothelial cell monolayers was performed after five days of culture of endothelial cells in apical transwell chamber. Cells were treated with DENV, ZIKV, WNV, JEV, and YFV NS1 proteins for 3 hours at 37°C. Purified DENV (1×10^5^ FFU), ZIKV (1×10^5^ FFU), or KUNV (1×10^6^ FFU) was then added to the apical side and incubated for 24 hours. The number of viral particles that crossed through endothelial cell monolayers were measured by RT-qPCR as described above. Virus dissemination through endothelial cells was expressed as log-fold changes of the viral GE/mL obtained under distinct experimental conditions normalized to the amount of virus detected in the basolateral chamber in untreated conditions. For both assay systems, TEER was measured 3 hours post-treatment with NS1 right before virus or U937 cells were added to the apical or basolateral sides of transwell inserts containing polarized endothelial cells (see TEER assay for endothelial permeability).

### Fluorescence microscopy

For immunofluorescence imaging experiments, normal human astrocytes (NHA) were grown on glass coverslips coated with 0.2% gelatin and imaged on a Zeiss LSM 710 Axio Observer inverted fluorescence microscope equipped with a 34-channel spectral detector. Briefly, NHA grown on coverslips were fixed using formaldehyde (FA 4%), and staining for NS3 viral protein using a primary mouse IgG mAb (E1D8) was performed after permeabilization followed by addition of a secondary species-specific anti-IgG antibody conjugated to a fluorophore (Alexa 647). Images acquired using the Zen 2010 software (Zeiss) were processed and analyzed with ImageJ software as previously described.^25^ Additionally, staining of NS3 was performed on endothelial cells directly grown on transwell inserts after the virus dissemination experiments. Briefly, after fixing the endothelial cell monolayer on the transwell using formaldehyde (FA 4%), the transwell was carefully removed from the plastic basket, and the staining was performed as described above. All RGB images were converted to grayscale, then mean grayscale values and integrated density from selected areas were taken along with adjacent background readings and plotted as mean fluorescence intensity (MFI). Images are representative of at least n=3 biological replicates.

### NS1-virus interaction assays

Purified NS1 proteins (DENV and ZIKV) with C-terminal His tags were diluted to 5 μg/mL in NTE buffer, then mixed with Ni-NTA beads (Thermo-Fisher) for 2 hours at 4°C while rotating end-over-end. NS1-bound beads were washed 2X with PBS, then purified virus (DENV or ZIKV) was added in NTE buffer and allowed to bind for 2 hours at 4°C while rotating. Beads were washed 2X with cold NTE buffer, then 2 with 30mM Imidazole in NTE buffer, and were then eluted with 300mM Imidazole. Eluted NS1 and virus were collected, and vRNA was purified and quantified as previously described above for RT-qPCR.

### Statistics

Prism version 5.0 (GraphPad) was used for data analysis. For statistical comparisons between 2 groups of data, unpaired nonparametric tests such as t test (Mann-Whitney) and one-way ANOVA (Kruskal-Wallis test) were used to determine the difference between medians inside different groups. For multiple comparison between groups, ordinary two-way ANOVA was used. A p-value of <0.05 was considered a statistically significant difference.

## Acknowledgements

We thank members of the Harris Laboratory at UC Berkeley, including Felix Pahmeier, Laurentia Tjang, and Richard Ruan for helpful discussion and technical assistance. Confocal imaging experiments were conducted on a Zeiss LSM 710 at the CRL Molecular Imaging Center, supported by the Gordon and Betty Moore Foundation. This work was supported by NIAID/NIH grants R01 AI168003 and R01 AI24493 (E.H.). S.B.B. was supported in part as an Open Philanthropy Awardee of the Life Sciences Research Foundation. The work was also supported by the Judith and Trent Anderson Endowment (R.J.K).

## Author Contributions

Conceptualization: HPG, SBB, PRB, EH

Data curation: HPG, SBB, RJK, EH

Formal analysis: HPG, SBB, MJDBW

Funding acquisition: RJK, EH

Investigation: HPG, SBB, BCR, MJDBW, NTNL, DAS, CMW, CW, TC, DRG

Methodology: HPG, SBB, MJDBW, RJK, EH

Project administration: HPG, SBB, RJK, EH

Resources: RJK, EH

Supervision: HPG, SBB, RJK, EH

Validation: HPG, SBB, MJDBW

Visualization: HPG, SBB, EH

Writing – original draft: SBB, HPG, EH

Writing – review & editing: HPG, SBB, RJK, EH

---

**S1 Fig.**
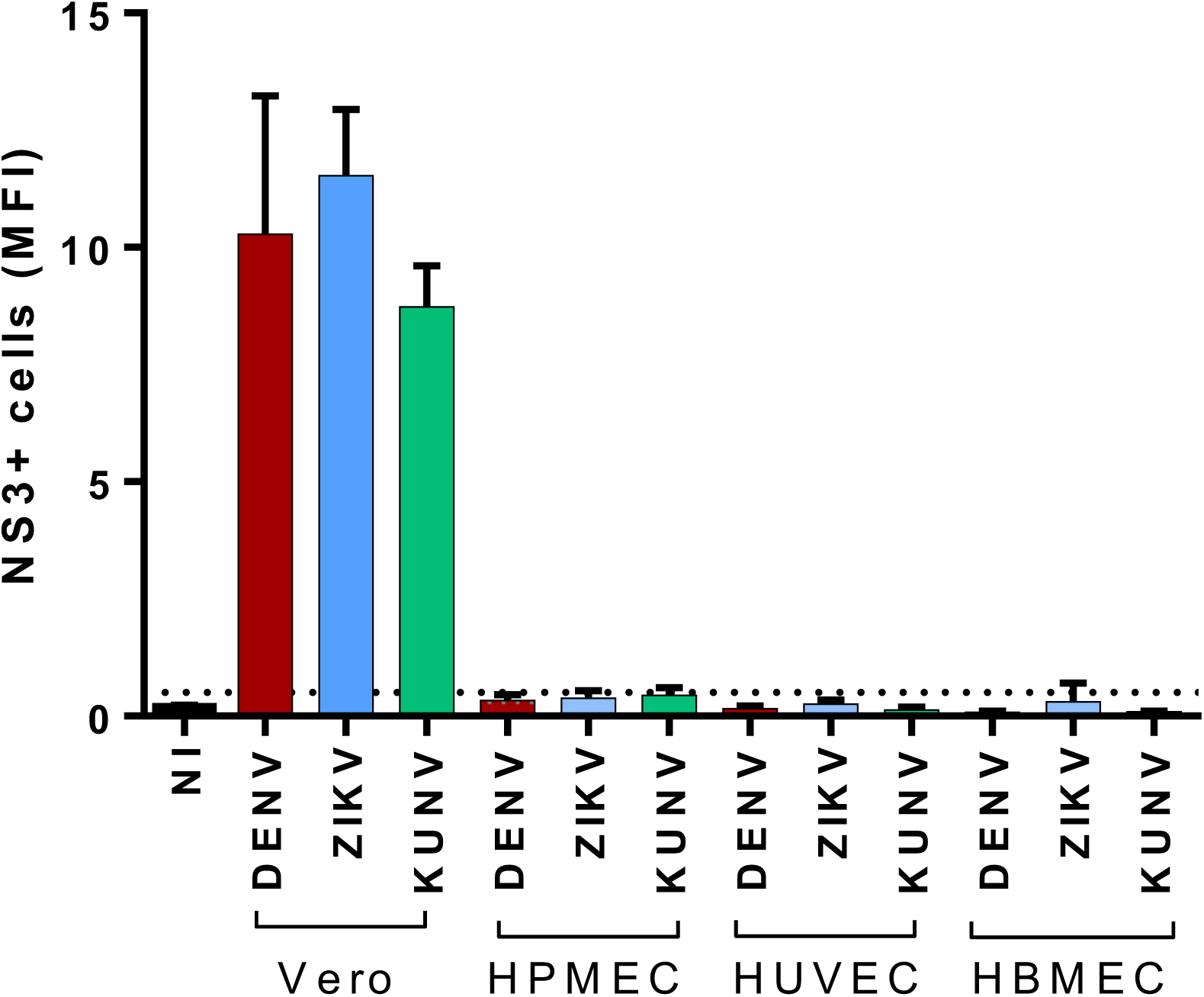
DENV, ZIKV, and KUNV do not infect endothelial cells *in vitro* at 24 hours post-infection. Monolayers of HPMEC, HUVEC, HBMEC, or Vero-CCL81 cells grown on transwells were treated for 3 hours with the indicated NS1 proteins (5 µg/mL). They were then treated with 1×10^5^ FFU of DENV, 1×10^5^ FFU of ZIKV, or 1×10^6^ FFU KUNV (added to the apical chamber) as in Figure 2 and 3. Twenty-four hours later, cells were fixed in the transwells, and viral infection was monitored by IFA staining for NS3. Quantification of mean fluorescence intensity (MFI) was normalized to cell number.

